# Effects of Host-Tree Foliage on Polymorphism in an Insect Pathogen

**DOI:** 10.1101/2024.09.29.615698

**Authors:** Ari S. Freedman, Amy Y. Huang, Katherine P. Dixon, Carlos Polivka, Greg Dwyer

**Author notes:** These authors contributed equally (Ari S. Freedman, Amy Y. Huang).

## Abstract

The theory of host-pathogen interactions has successfully shown that persistent pathogen virulence may be explained through tradeoffs between different pathogen fitness components, but classical theory cannot explain pathogen coexistence. More recent theory invokes both tradeoffs and environmental heterogeneity, but resembles classical theory in focusing on a limited range of possible tradeoffs, and therefore has seen few applications. To better understand the usefulness of tradeoff theory for explaining pathogen coexistence in nature, we measured components of pathogen fitness in two distantly related morphotypes of a baculovirus that infects larvae of the Douglas-fir tussock moth (*Orgyia pseudotsugata*). We show that the two morphotypes vary in multiple components of fitness, including the probability of infection given exposure to the pathogen, the incubation time of the pathogen, variability in the incubation time of the pathogen, and the detectability of the pathogen. Moreover, because the baculovirus is transmitted when host larvae accidentally consume infectious virus particles while feeding on foliage of the insect’s host trees, the strength and direction of the differences in fitness components of the two morphotypes depends on the host-tree species on which host larvae consume the virus. Through simulations of a model parameterized using our experimental data, we demonstrate how several varying fitness components can work in concert to promote strain coexistence, particularly highlighting the role of variability in incubation time. Our results suggest that the two morphotypes may coexist because of variation in forest tree-species composition, providing important empirical evidence that tradeoffs and environmental heterogeneity can together modulate pathogen competition.

## Introduction

Pathogen polymorphism is ubiquitous and widely observed (Buckee et al., 2007; Domingo et al., 1985; Hitchman et al., 2007; Lipsitch and O’Hagan, 2007), an observation that long posed a difficult challenge for classical theory of host-pathogen interactions. Classical theory emphasized trade-offs in pathogen fitness components, in an effort to understand the evolution of intermediate virulence (Anderson and May, 1982). Simple tradeoff theory, however, predicts that fitter strains should always competitively exclude less-fit strains (Keeling and Rohani, 2008), preventing polymorphism.

More complex models allow for pathogen coexistence through mechanisms that include spatial structure (Messinger and Ostling, 2009), seasonality (Andreasen and Dwyer, 2022), and heterogeneity in host or pathogen population structure (Fleming-Davies et al., 2015). Even these more complex models, however, oversimplify the ways in which pathogen fitness may differ in different environments. To better understand the ways in which environmental heterogeneity may affect pathogen competition, here we study mechanisms that affect the fitness of two competing morphotypes of a baculovirus of the Douglas-fir tussock moth, *Orgyia pseudotsugata*.

Like other baculoviruses, the tussock moth baculovirus is an obligately-lethal pathogen, meaning that it must kill its host before it can be transmitted (Dwyer, 1992). Transmission occurs when uninfected larvae consume occlusion bodies, virion-containing protein matrices that are released onto foliage from virus-killed cadavers (Federici, 1997). The tussock moth baculovirus consists of two morphotypes that differ in the ultrastructure of their occlusion bodies (Hughes and Addison, 1970); previous work provided modest evidence that the two morphotypes differ in their speed of kill (Hughes, 1976), but little other information.

The two morphotypes have been shown to co-occur at high frequency at many locations across the range of the insect (Williams et al., 2011), including a large fraction of the western USA and the province of British Columbia, Canada, and data demonstrating co-occurrence have been collected over multiple decades (Hughes, 1976; Hughes and Addison, 1970). Previous data therefore strongly suggest that the two morphotypes coexist, and in what follows we investigate potential mechanisms to explain this apparent coexistence.

To identify mechanisms that may explain the apparent coexistence of the two morphotypes of the tussock moth baculovirus, we quantified key components of the fitness of each morphotype. Lepidopteran-baculovirus interactions are ideal for breaking down pathogen fitness into its underlying components. The pathogen’s transmission pathway is relatively simple yet hosts often exhibit a high degree of heterogeneity in infection risk (Dwyer et al., 1997; Mihaljevic et al., 2020), suggesting that transmission is more complex than assumed by many mathematical models. Moreover, because successful infections invariably lead to host death, it is possible to determine the exposure period by recording when a host larva contacts the pathogen by consuming contaminated foliage and when such contact leads to a successful infection by recording when the larva dies of infection.

Mathematical models of host-pathogen interactions generally describe transmission using a single parameter (Jagan et al., 2020; Kirkeby et al., 2017), following a mass-action assumption in which transmission depends only on the product of the densities of susceptible and infected hosts (McCallum et al., 2001). Transmission in nature by contrast often consists of numerous sub-processes, and so empiricists have called for efforts to unpack the biological processes contributing to transmission to better inform modelling efforts (Antolin, 2008; LaDeau et al., 2011). Following these calls, we unpacked the transmission of the tussock moth baculovirus into two general components, the probability of death given consumption of the virus—a measure of lethality—and the probability that the larva consumes the virus in the first place, a measure of exposure risk.

Because theories of host-pathogen competition have further identified incubation times as key components of pathogen fitness (Anderson and May, 1982), and because previous work suggested that incubation times may differ between the two morphotypes, we also quantified the speed of kill of each morphotype. Although previous work on the fitness of obligately-lethal pathogens considered only mean speed of kill (Ebert and Weisser, 1997), we further measured variance in the speed of kill, which as we show can have important effects on pathogen fitness and promote polymorphism.

An important feature of the biology of the Douglas-fir tussock moth is that larvae can successfully complete their development on both the eponymous Douglas-fir (*Pseudotsuga menziesii*) and on multiple species of so-called “true firs” in the genus *Abies* (Wickman et al., 1981). This is important for the coexistence of the two morphotypes of the tussock moth baculovirus, as previous work with baculoviruses of other insects has shown that the probability of death given virus consumption can vary between host-plant species (Dwyer et al., 2005; Elderd et al., 2013), and that speed of kill can vary with viral strain and host-plant species in several other baculovirus systems (Ali et al., 2002; Duffey et al., 1995; Hodgson et al., 2002; Hoover et al., 1998; Keating et al., 1988; Shikano et al., 2017). Interacting effects of viral strain and host-plant species, however, are poorly understood (Hodgson et al., 2002). Similarly, although pathogen-avoidance behavior has been documented in larvae of the closely-related spongy moth (*Lymantria dispar*) (Capinera et al., 1976; Eakin et al., 2015; Parker et al., 2010), the possibility that host-plant foliage or viral strain may affect avoidance ability has not been considered. To test whether differences in fitness components between viral morphotypes may depend on differences in host-plant foliage, we therefore quantified the fitness of each viral morphotype on Douglas-fir and grand fir, *Abies grandis*.

Our work shows first that host-tree species, viral morphotype, and interactions between host-tree species and viral morphotype have strong affects on the virus’s speed of kill, on the risk of death given virus consumption, and on the larva’s ability to avoid virus consumption in the first place. We further show that host-tree species, viral morphotype, and interactions between host-tree species and viral morphotype affect the baculovirus’s mean speed of kill and variance in the speed of kill; using simulations of an SEIR (susceptible-exposed-infected-recovered) disease model, we show that strains with higher variation in their speed of kill have higher fitness because a greater proportion of their kills occur in a shorter period of time. This latter result represents one of the first theoretical demonstrations that differences in the variance of a pathogen’s incubation time can alter pathogen fitness (a few previous efforts have come from plant disease modeling; Ferrandino 2012, 2013; Suffert and Thompson 2018).

Because the relative frequency of Douglas-fir and true firs varies across the range of the insect, our work shows that the transmission dynamics of the tussock moth baculovirus and the competitive balance between the viral morphotypes both depend on variation in forest composition. Our work thus provides insight into mechanisms that facilitate pathogen coexistence, and demonstrates the conceptual usefulness of teasing apart pathogen fitness into its component parts.

## Methods

As we described, we focused on two components of baculovirus transmission: (1) risk of exposure and (2) risk of infection given exposure. To measure exposure risk, we used choice tests in which larvae were allowed to choose between virus-contaminated foliage and uncontaminated foliage. This experiment used a modification of a protocol first developed for the spongy moth, which can detect and avoid virus-contaminated foliage while feeding near infectious cadavers (Eakin et al., 2015; Hudson et al., 2016). We quantified risk of infection given exposure by feeding the larvae contaminated foliage and recording the fraction that became infected and therefore died. Note that because the virus is obligately lethal, only infections that end in death are relevant to pathogen fitness.

We also considered two components of the pathogen’s incubation time that affect pathogen fitness: the mean speed of kill and the variance in the speed of kill. We again measured these fitness components using infection experiments in which we fed virus-contaminated needles to larvae and discarded larvae that did not consume an entire needle. To understand the consequences of variation in the mean speed of kill and the variance in the speed of kill for pathogen fitness, we inserted our measurements into a simple host-pathogen model and we used the model to project the effects of the average and the variance in the speed of kill on pathogen fitness and competitiveness.

All of the components of baculovirus transmission that we measure and the methods used to measure them are summarized in Fig. 1.

**Figure 1:**
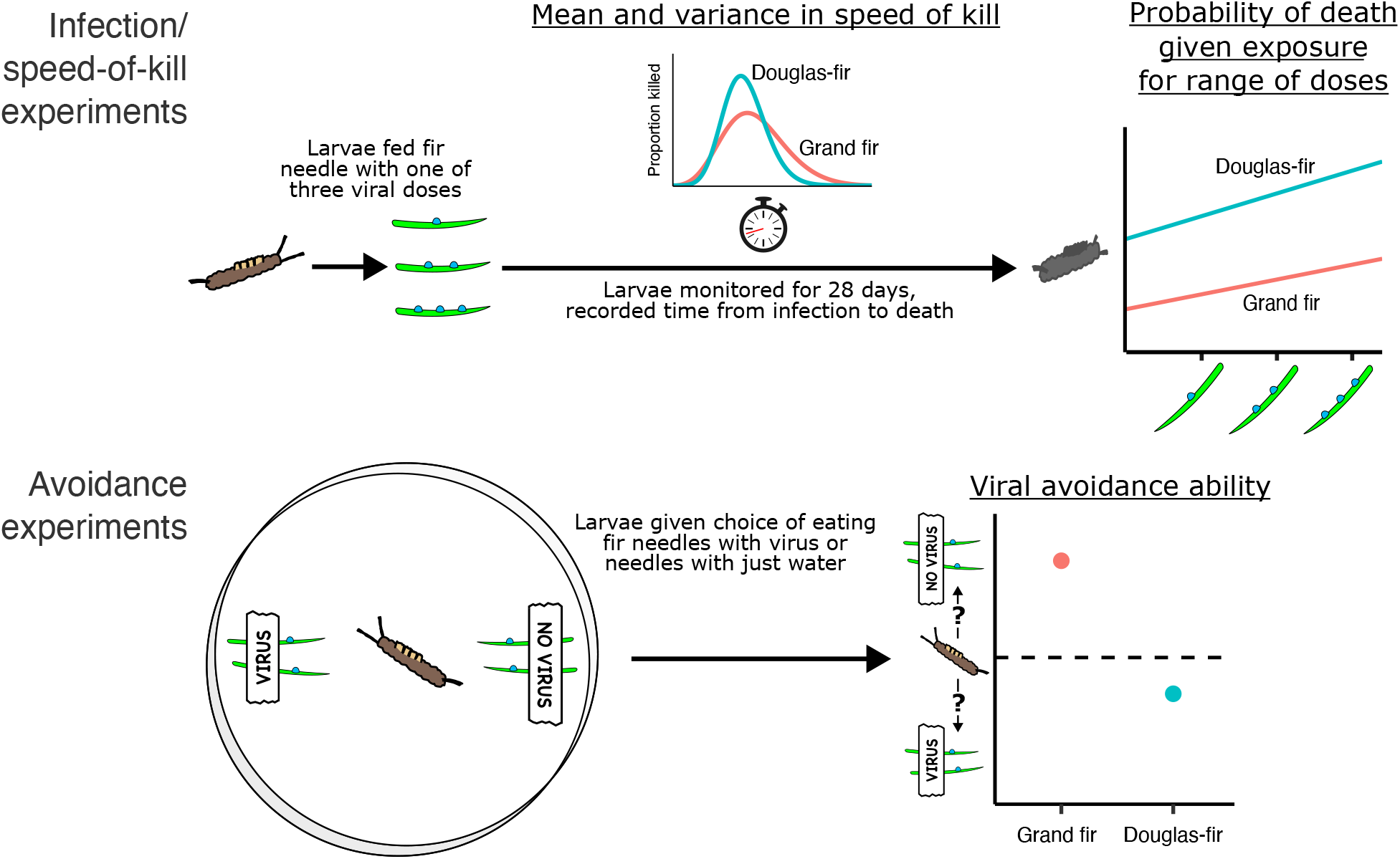
Conceptual depiction of our experimental methods. In infection experiments, we fed larvae with one of three viral doses in water drops deployed on top of fir needles. We then recorded the fraction of larvae killed, as well as the mean and variance in the time it took for infected larvae to die. In avoidance experiments, we gave larvae a choice between fir needles with virus solution and fir needles with plain water, and we recorded the needle area before and after larval feeding. Fir needles were taken from the field from branches of either grand fir or Douglas-fir, and we used several different isolates of each of the two morphotypes of the baculovirus, SNPV and MNPV. The pathogen fitness components we measured are underlined.

### Natural history of the Douglas-fir tussock moth and its baculovirus

The Douglas-fir tussock moth is an economically important native defoliator of Douglas-fir and true firs in the genus *Abies* in Western North America, and its periodic outbreaks result in extensive defoliation (Beckwith, 1976). In the northwestern United States, where our experiments were conducted, larvae hatch in late May from eggs laid the previous August/September. Tussock moth outbreaks last for 2–4 years and are typically ended as a result of the environmentally-transmitted baculovirus, specifically a nucleopolyhedrovirus or “NPV” (Mihaljevic et al., 2020; Otvos et al., 1987). Epizootics begin when neonates that have hatched from virus-contaminated eggs die of the virus, serving as a source of virus that infects additional larvae (Dwyer and Elkinton, 1993). Subsequent infections come from consumption of foliage contaminated by virus-killed cadavers; thus, host diet and foraging intensity are important determinants of virus exposure.

The nucleopolyhedrovirus consists of two morphotypes, which differ in how their virions are packaged within the infectious particles or “occlusion bodies”. In the single-capsid (“SNPV”) morphotype virions or “capsids” occur singly within occlusion bodies, while in the multi-capsid (“MNPV”) morphotype virions occur in bundles within the occlusion bodies (Hughes and Addison, 1970; Rohrmann et al., 1978). Surveys of morphotype frequencies have observed high levels of both morphotypes within local populations, with the dominant morphotype varying between populations (Dixon, 2024; Shepherd et al., 1984; Williams et al., 2011).

### Infection experiments

To measure the risk of infection given exposure, we starved larvae for 48 hours, fed the larvae virus-contaminated needles, and discarded larvae that did not consume an entire needle. Because baculovirus infection rates are strongly affected by variation in virus doses and host-tree foliage quality (Dwyer et al., 2005), in our experiments we varied both the tree species from which we obtained needles for our experiments and the virus dose that we fed to the larvae. The needles came from Douglas-fir and grand fir branches that we obtained from mixed Douglas-fir/grand-fir forests near Beehive Reservoir outside of Wenatchee, Washington (47.330^°^N, −120.403^°^W). The virus doses consisted of 1,800, 3,600, and 5,400 occlusion bodies, delivered in 1, 2, or 3 *µ*L drops of 1,800 occlusion body/*µ*L virus solution, as 3 *µ*L was the maximum amount of liquid we were able to fit onto a single fir needle. Because it is not possible to exactly reproduce a given dose, the density of occlusion bodies varied slightly between the solutions we produced for each isolate (Table S1 in Online Supplement).

Although previous studies have provided some evidence that phenotypes vary between isolates in other insect-baculovirus-insect systems (Cory et al., 1997; Fleming-Davies et al., 2015), little is known about variation in phenotype across isolates of the two morphotypes of the tussock moth virus. We therefore used eight isolates collected from sites distributed across the western USA, such that four were SNPV, labeled as COL, SUB, LOV, and LST, while four were MNPV, labeled as DRY, KLP, TAM, TMB (Table S1). The TMB was a sample of the bio-pesticide “TM Biocontrol-1” isolate (Martignoni, 1999).

Virus solutions were pipetted directly onto single fir needles, and each larva was fed one needle after the liquid had evaporated. After 36 hours, larvae that did not consume their entire needle were discarded, to ensure that all larvae ingested the entirety of their viral doses. Control larvae were fed needles with water only as we described.

A total of 653 larvae consumed needles with SNPV isolates, 666 larvae consumed needles with MNPV isolates, and 74 control larvae consumed needles with distilled water (see Table S2 for breakdown by isoalte and tree species). Only 1 out the 74 control larvae (1.4%) became infected, a contamination rate that is low enough that we do not consider control larvae in what follows. We reared larvae on artificial diet for a total of 28 days post-infection, a period long enough to ensure that all infected larvae would die of the infection (Kennedy et al., 2014). To quantify speeds of kill, we recorded mortality each day starting six days after infection, the minimum time needed for the virus to kill its host from first consumption (Morris, 1963). Because occlusion bodies are clearly visible at 400*×* magnification under a light microscope, dead larvae were autopsied with a compound microscope to confirm deaths owing to the virus.

### Baculovirus avoidance in choice tests

To quantify the ability of larvae to avoid foliage contaminated with infectious cadavers, we carried out choice tests. For each larva in the choice test, we placed two needles of uncontaminated foliage and two needles of contaminated foliage in a Petri dish, such that needles were taken from either grand fir or Douglas-fir branches. Because the neonate larvae that were infected for transmission naturally select foliage derived from the same year’s new buds, we used that season’s new foliage. We tested four viral isolates, two SNPV and two MNPV. Control plates were identical to treatment plates but contained uncontaminated needles. We carried out this procedure for 63 larvae with SNPV isolates, 62 larvae with MNPV isolates, and 36 control larvae.

One of the most important rounds of baculovirus infection in nature occurs when tussock moth larvae in the fourth larval stage or “instar” consume needles contaminated with the cadavers of neonates (Mihaljevic et al., 2020). To replicate natural conditions as closely as possible, for the choice tests we produced cadaver-contaminated needles by allowing neonates to die naturally on foliage of grand fir or Douglas-fir branches in the field, again using trees at Beehive Reservoir. To do this, we placed infected neonates on branches enclosed in mesh bags, and allowed them to die. To ensure that the neonates were infected, we used a dose 2,000 occlusion bodies per *µ*L which was sufficient to ensure 100% infection. To ensure that all neonates died on the branches, we left them in the field for 13 days (infected neonates die faster than infected fourth instars). Approximately 75 infected larvae were placed in each bag. To ensure that conditions for the control needles mirrored the conditions of the contaminated foliage, we collected control needles from branches that were similarly enclosed in mesh bags but that contained no infected larvae.

Because cadavers are clearly visible on needles, we were easily able to select cadaver-contaminated needles for the experiment. We then placed these needles in Petri dishes, added larvae in the fourth instar, and allowed each larva to feed for 24 hours. Each Petri dish contained fir needles from just one tree species, either grand fir or Douglas-fir. To quantify the amount of contaminated and uncontaminated needles that each larva consumed, we photographed the needles before and after larval feeding. To quantify the difference in needle area before and after feeding, and thus to quantify the amount of virus-contaminated needle that each larva consumed, we used the software app “ImageJ” to measure needle areas (Schneider et al., 2012). We then used the difference in the amount of contaminated versus uncontaminated foliage eaten by each larva as a measure of cadaver avoidance.

### Statistical analysis and model fitting

Because isolates within a morphotype share a common descent and are thus not independent (Rohrmann et al., 1978), we used Bayesian hierarchical models to fit statistical models to our dose-response and avoidance data. This meant that we simultaneously fit model parameters separately by isolate and morphotype-level parameters that determined the distributions from which the isolate-level parameters are drawn. This procedure allowed us to take into account both differences between isolates within a morphotype and differences between the morphotypes themselves. Because each individual isolate did not have enough data points to provide reliable estimates of an isolate-specific speed-of-kill distribution, and because there was no obvious way to construct a hierarchical model for our speed-of-kill data, our analyses of speed of kill combined the data from all isolates within a morphotype.

For all of our analyses, we chose the best model using the leave-one-out cross-validation or “LOO-CV” information criterion. LOO-CV is a Bayesian analog to the traditional Akaike information criterion or “AIC” that is especially suitable for small data sets (Vehtari et al., 2016). Our models were implemented in the statistical programming language Stan (Stan Development Team, 2024).

Our dose-response models allowed us to test for effects of morphotype and tree species on the probability of death given exposure. We constructed four different versions of these models (Table 1), such that each model followed the general “logit” form:

**Table 1:** LOO-CV analysis of our dose-response models. For each model we show the LOO estimate of the expected log pointwise predictive densities (ELPD), the difference in ELPD from the best model (ΔELPD), and the standard errors of these differences (SE). The best model has ΔELPD = 0 and worse-fitting models have larger negative values of ΔELPD.

| Model | ELPD | ΔELPD | SE |
| --- | --- | --- | --- |
| Morphotype + tree species | -115.0 | 0.0 | 0.0 |
| Tree species only | -117.6 | -2.6 | 1.7 |
| Morphotype only | -223.6 | -108.6 | 23.0 |
| Intercept | -227.2 | -112.2 | 22.7 |

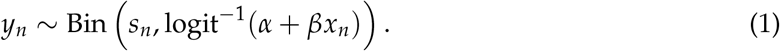

Here, Bin(*k, p*) is a binomially-distributed random variate with *k* trials and probability *p* of success. Larvae in treatment *n* have probability of death given exposure *y*_*n*_, sample size *s*_*n*_, and dose *x*_*n*_ in units of occlusion bodies; *α* and *β* are fitted parameters; and logit^−1^ : ℝ → (0, 1) is the inverse-logit or logistic function defined as logit^−1^(*x*) = 1/(1 + *e*^−*x*^). In all models, *α* and *β* differed between isolates. In models that include morphotype as a factor, *α* and *β* were drawn from different distributions depending on the isolate’s morphotype. In models that included tree species as a factor, *α* and *β* and their associated morphotype-level parameters also varied by tree species. To test the assumption of the isolate-morphotype hierarchy, we also included a model variant in which isolate and tree species were factors but for which there was no morphotype-based hierarchy (see Online Supplement). Because as we will show some dose-response curves had near-zero slopes, in all dose-response models we constrained *β* to be non-negative to avoid the possibility of unrealistic (small) negative slopes of the dose-response curves.

To determine if morphotype and tree species affected the virus’s speed of kill in our infection experiments, we analyzed our speed-of-kill data using gamma distributions. Gamma distributions are widely used in epidemiological models to describe incubation times (Keeling and Rohani, 2008), including speeds of kill (Mihaljevic et al., 2020). The underlying assumption is that an infected individual passes through a series of “exposed” classes, such that the time in each class is exponentially distributed. The sum of the times in all the classes together, and thus the speed of kill, then follows a gamma distribution. Because visual inspection of the data made clear that the doses that we used had no more than minor effects on speeds of kill, for the purposes of fitting speed-of-kill distributions we pooled larvae across doses and isolates within a morphotype. We then allowed the parameters of our gamma distributions to differ across morphotype, tree species, both morphotype and tree species, or neither morphotype or tree species, to produce a total of four models.

In the choice tests, we quantified the degree of avoidance by measuring the difference in the percent eaten over a 24-hour period between the contaminated and the uncontaminated leaves.

We symbolize this avoidance metric as *D*_*n*_, defined as:

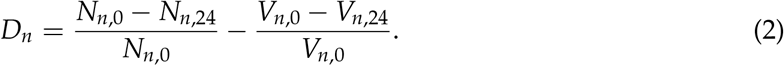

Here *N*_*n,t*_ is the total remaining leaf area of the “no virus” side of plate *n* after *t* hours of feeding, while *V*_*n,t*_ is the remaining leaf area of the “virus” side of plate *n* after *t* hours. In the no-virus control dishes, neither side contained virus-contaminated needles, but we nevertheless labeled the sides labeled “virus” and “no virus” for consistency during imaging and bias correction.

Because we were concerned that our measurements of the area consumed would be sensitive to variation in light during the photographic process, we corrected our *D*_*n*_ metric by by subtracting off the average *D*_*n*_ as measured from the control plates. Our bias-corrected avoidance metric, 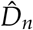 is then calculated as:

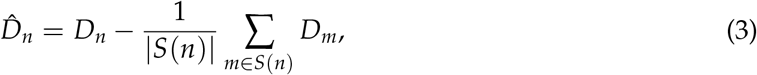

where *S*(*n*) is the set of control plates with the same tree species as in plate *n*, and |*S*(*n*)| is the size of *S*(*n*), and thus the number of control plates that have the same tree species as plate *n*. In other words, 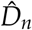 is equal to *D*_*n*_ minus the average value of *D*_*m*_ for control plates *m* of the same tree species. A positive value of 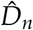 means the larva in plate *n* tended to avoid virus-contaminated foliage, a negative value means the larva preferred contaminated foliage, and 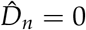 means that the larva did not discriminate between contaminated and uncontaminated foliage.

Our avoidance models quantified the relative importance of viral presence, morphotype, tree species, and morphotype-tree species interaction. Our most complex avoidance model includes all of these factors:

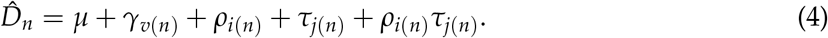

Here 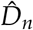 is the model’s estimate of the avoidance metric for Petri dish *n*; *γ*_*v*(*n*)_ accounts for viral presence, where *v*(*n*) represents whether *n* is a treatment or control plate, and *γ*_control_ = 0; *ρ*_*i*(*n*)_ accounts for viral morphotype and isolate, such that *i*(*n*) is the isolate used in plate *n* but the standard deviation of *ρ* varies only by morphotype, and *ρ*_control_ = 0; and *τ*_*j*(*n*)_ accounts for tree species, such that *j*(*n*) is the tree species used in plate *n*.

We used Markov chain Monte Carlo (MCMC) to fit our models to data, such that dose-response and speed-of-kill models were run with 4 chains of 4,000 iterations each and avoidance models with 5 chains of 10,000 iterations each. We used uninformative priors for each parameter in all models (see Online Supplement).

For each model, we calculated the LOO estimate of the expected log pointwise predictive density or “ELPD”, which is equal to minus one-half the LOO-CV value, the difference in ELPD between each model and the best model for each data set, and the standard errors of these ELPD differences. The best model then has the highest ELPD. Following standard practice (Sivula et al., 2020; Vehtari et al., 2016), we concluded that the best model provides a meaningfully better explanation for a data set if the magnitude of the difference in ELPD between the best model and any other model fit to that set exceeded the standard error of the ELPD difference.

### Understanding the consequences of variation in speed of kill for pathogen fitness

As we described earlier, the theoretical epidemiology literature has focused on mean incubation times or mean speeds of kill to the exclusion of the variance in the incubation time or speed of kill (Ebert and Weisser, 1997). We therefore know little about how differences in the variance in the incubation time affects pathogen competitive ability. Although the same is true of variation in cadaver avoidance behavior and of variation in the dose required to cause infection, models that allow for variation in cadaver avoidance and for variation in the infectious dose can only be implemented using highly complex computer algorithms (Eakin et al., 2015). Although constructing such a model was beyond what we could accomplish, constructing a model that allows for variation in the variance in the speed of kill required only that we numerically integrate an SEIR model, which is comparatively straightforward.

To understand the implications of our speed-of-kill data for pathogen competitive ability, we therefore inserted our estimates of the mean and variance in speed of kill into an SEIR model that had previously been fit to data on baculovirus epizootics in Douglas-fir tussock moth populations (Mihaljevic et al., 2020). That previous work, however, assumed that the variance in the speed of kill was near zero; as we will show, however, the variance in the speed of kill of the tussock moth baculovirus is modest but certainly not zero. We therefore used the parameter estimates of the SEIR model from the previous work, except that we substituted our experimental estimates of the mean and the variance in the speed of kill.

The SEIR model is (Fuller et al., 2012):

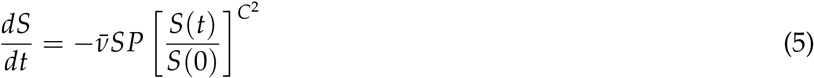

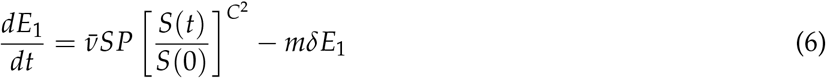

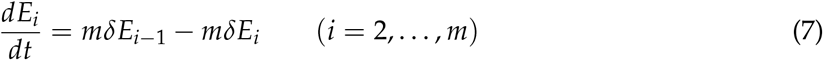

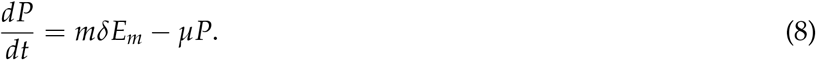

Here *S*(*t*) is the density of susceptible hosts at time *t, P*(*t*) is the density of infectious cadavers, and *E*_*i*_(*t*) for *i* = 1, …, *m* is the density of exposed but not yet infectious hosts. By including multiple exposed classes, we can allow for gamma-distributed speeds of kill, following the so-called “linear chain trick” (Smith, 2011). This is a standard approach in theoretical epidemiology, because it makes it possible to represent a distributed delay using ordinary differential equations, thereby considerably simplifying the computational overhead required to numerically integrate the model (Keeling and Rohani, 2008). The mean speed of kill is then 1/*δ* while the variance is 1/(*mδ*^2^). We then used our estimates of the mean and the variance in the speed of kill from our experimental data to estimate *δ* and *m*. In practice, this meant that we rounded *m* to the nearest integer, but the amount of round-off involved was quite small compared to the size of *m*, which ranged from 7 to 28. See the Online Supplement for more information on the parameter values used.

We simulated the transmission model using a period of 50 days, which is roughly the length of the epizootic during the larval stage. We began the model epizootics with a small amount of initial pathogen density *P*(0) = .01 cadavers/m^2^, as epizootics tend to start with a low initial density of virus on foliage left over from the previous season (Thompson and Scott, 1979). Because baculovirus fitness effects can vary with host density (Fleming-Davies et al., 2015), we allowed host densities to range from from .5 to 100 larvae/m^2^.

As a measure of pathogen fitness we used the cumulative fraction infected during a single epizootic, thereby following previous theory for strongly seasonal host-pathogen interactions (Andreasen and Dwyer, 2023). We then compared fitness outcomes between morphotypes on different tree species under three scenarios: (1) a scenario in which each of the four morphotype-tree species combinations has its own distinct, speed-of-kill distribution taken from our best-fit model; (2) a scenario in which each morphotype-tree species combination has its own distinct variance in speed of kill, but the two morphotypes have the same mean speed of kill, which is equal to the average speed of kill across all morphotypes and tree species; and (3) a scenario in which each morphotype-tree species combination has its own distinct mean speed of kill, but they all use the same variance in speed of kill. We present simulations from the first two scenarios here in the main text, and the third scenario in the Online Supplement.

## Results

### Dose response and the probability of infection given exposure

Larvae feeding on Douglas-fir needles had a higher probability of successful infection after exposure than larvae feeding on grand fir, with 43.6% virus infected on Douglas-fir and 19.3% on grand fir. Similarly, MNPV isolates tended to be more infectious than SNPV isolates, with 45.9% virus infected on MNPV and 17.6% on SNPV. Notably, however, the difference in infection rates between the two tree species was much more pronounced for MNPV isolates (66.0% on Douglas-fir, 26.9% on grand fir) than for SNPV (23.4% on Douglas-fir, 10.5% on grand fir; Fig. 2a).

**Figure 2:**
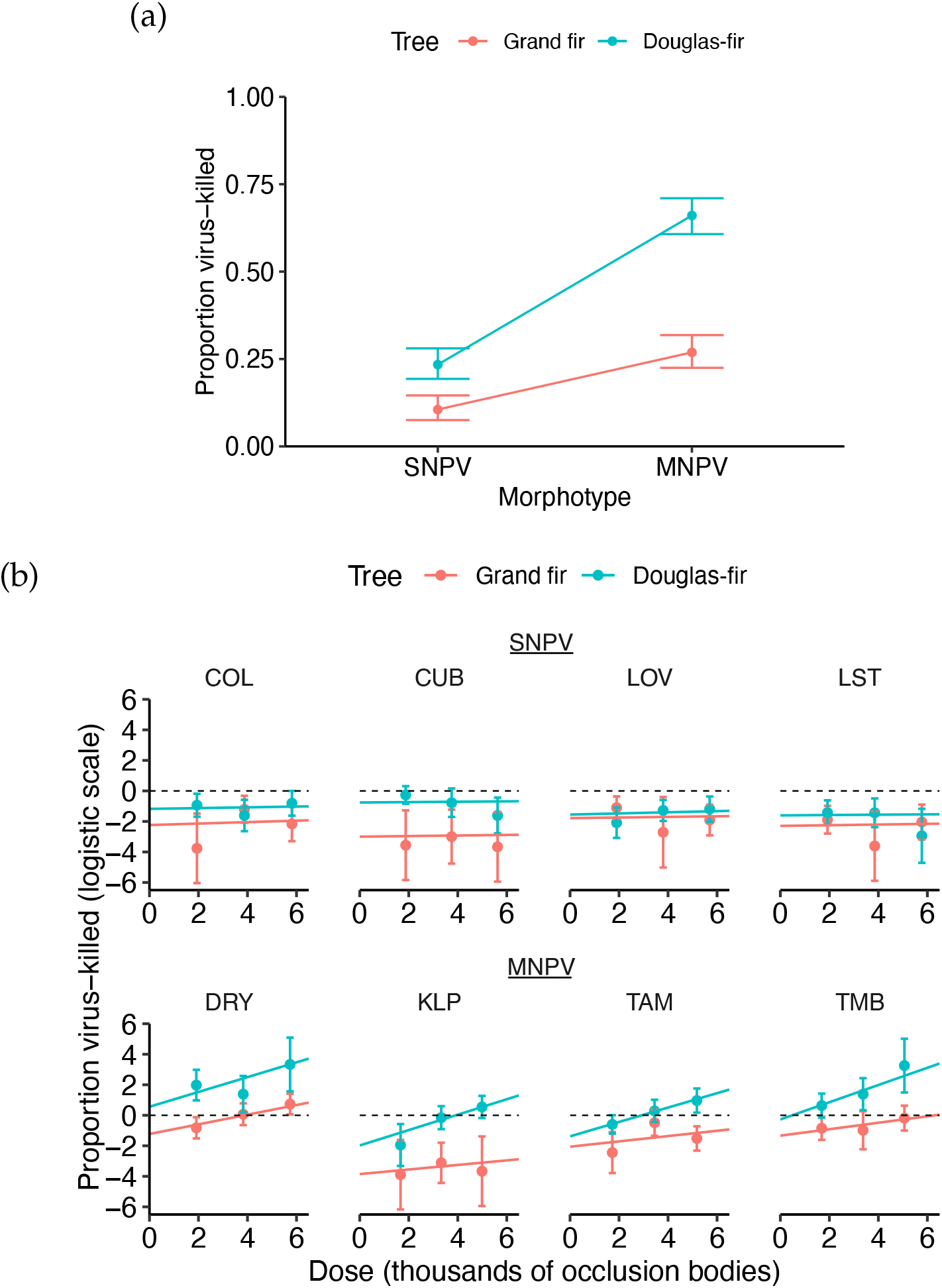
(a) Fraction of larvae killed by the tussock moth baculovirus in our infection experiments. (b) Fraction infected versus dose across isolates and tree species. Points represent fraction infected on a logit scale, while the lines show the fit of the best logit model, which includes effects of both morphotype and tree species. The dashed lines at 0 represent 50% lethality (LD50). To avoid taking the log of zero fractions, for the data points in treatment groups with no infected larva, we assumed that a doubled sample size would lead to one virus-killed larva. We used this approach only to visually represent the data, whereas in our statistical analyses we used the unchanged data. Error bars in both plots represent 95% binomial confidence intervals.

Confirming these trends, the best dose-response model as identified by LOO-CV allowed for effects of both morphotype and tree species, whereas models that did not allow for morphotype or tree species provided significantly worse explanations for the data (Table 1). Reassuringly, the best model does a good job of predicting the fraction infected in most treatments (Fig. 2b). The model that allows for tree species only (ΔELPD = −2.6, SE = 1.7) performs far better than the morphotype-only model (ΔELPD = −108.6, SE = 23.0), emphasizing the importance of tree species in determining baculovirus fitness.

Our estimates for the slope parameter *β* in the best dose-response model were all strongly positive for the MNPV isolates but were effectively zero for the SNPV isolates (Fig. 2b). The effect of these near-zero slopes can be seen in the predicted increase of percent killed over the range of doses; from the low dose of 1,600 viral occlusions bodies to the high dose of 5,800 occlusion bodies, the best model predicts that the increase in the probability of death among SNPV isolates is only 1.3% on average (min. 0.4%, max. 2.5%), while the increase in the probability of death among MNPV isolates is 25.5% on average (min. 2.2%, max. 48.0%). The infectiousness of the SNPV morphotype was thus almost the same, and relatively low across virus doses.

### Cadaver avoidance and the probability of exposure

The avoidance experiments showed that tussock moth feeding behavior is sharply altered by the presence of virus-infected cadavers, but whether larvae avoided or sought out virus-contaminated needles varied strongly with morphotype and host-tree species. Overall, larvae that had a choice of cadaver-contaminated and uncontaminated foliage consumed slightly more uncontaminated foliage (mean 34.5%, SE 1.3%) than cadaver-contaminated foliage (mean 31.1%, SE 1.3%; *t*-test, *t*(319.3) = 1.83, *p* = .03). And as one would expect, larvae that were fed only uncontaminated foliage did not differ significantly in their foliage consumption between the needles on the sides marked as “virus” (mean 31.2%, SE 2.5%) and “no virus” (mean 30.1%, SE 3.1%; *t*(67.5) = .27, *p* = .79), but for completeness we report values of the avoidance metric 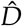 that take into account this slight bias.

Surprisingly, larvae feeding on Douglas-fir needles had a slight preference for needles contaminated with SNPV-infected cadavers (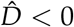, *t*(46) = 1.52, *p* = .07; recall that negative values indicate a preference for contaminated foliage over uncontaminated foliage), whereas larvae feeding on grand-fir needles had a non-significant preference for uncontaminated foliage 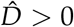, *t*(51) = 1.31, *p* = .10; Fig. 4a). In contrast, larvae strongly avoided needles contaminated with cadavers infected with MNPV isolates on both tree species (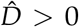, *t*(30) = 3.23, *p* < 0.01 on Douglas-fir; 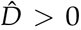, *t*(30) = 4.41, *p* < 0.01 on grand fir). Larvae were thus deterred more by MNPV-infected cadavers than by SNPV-infected cadavers, and deterrence was stronger on grand fir than on Douglas-fir.

Significance tests on larval avoidance ability thus showed that there were significant effects of viral morphotype and tree species. These results were confirmed by the results from our LOO-CV analysis of the avoidance models (Table 2), for which the best model accounted for viral presence, viral morphotype, and tree species, as well as the morphotype-tree species interaction. The best model produced a nearly perfect fit to the mean 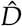 for each group (Fig. 4b). The next best model, which lacks the interaction term but is otherwise identical to the best model, performs significantly worse (ΔELPD = −5.0, SE = 4.4). Cadaver presence thus had strong effects on larval feeding behavior, as did viral morphotype, tree species, and the interaction between morphotype and tree species. The model with cadaver presence and tree species (ΔELPD = −24.9, SE = 8.5) provided a much worse fit to the data than did the model with cadaver presence and morphotype (ΔELPD = −7.9, SE = 5.2), demonstrating that morphotype is a better predictor than tree species of the effects of cadaver presence on larval feeding behavior.

**Table 2:**
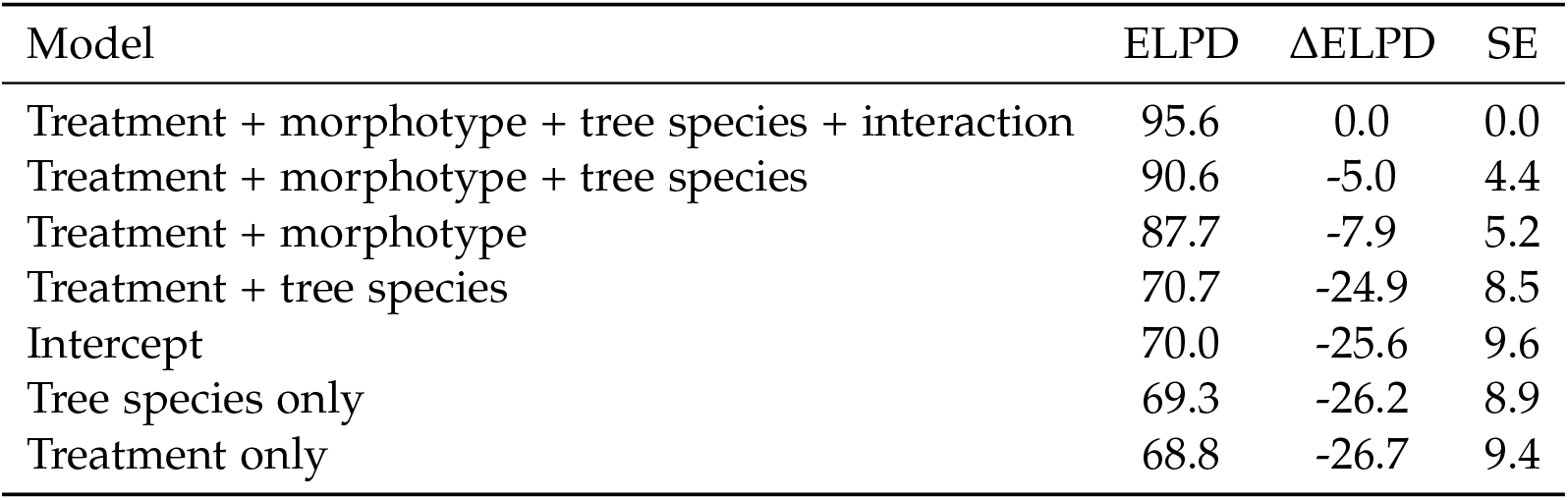
LOO-CV analysis of the cadaver avoidance models. For each model we show the LOO estimate of the expected log pointwise predictive density (ELPD), the differences in ELPD from the best model (ΔELPD), and the standard error of the difference (SE).

| Model | ELPD | $\Delta\text{ELPD}$ | SE |
| --- | --- | --- | --- |
| Treatment + morphotype + tree species + interaction | 95.6 | 0.0 | 0.0 |
| Treatment + morphotype + tree species | 90.6 | -5.0 | 4.4 |
| Treatment + morphotype | 87.7 | -7.9 | 5.2 |
| Treatment + tree species | 70.7 | -24.9 | 8.5 |
| Intercept | 70.0 | -25.6 | 9.6 |
| Tree species only | 69.3 | -26.2 | 8.9 |
| Treatment only | 68.8 | -26.7 | 9.4 |

### Speed of kill

#### Empirical results

For the MNPV morphotype, the speed of kill was significantly shorter on Douglas-fir (mean 12.7, SE .2 days) than on grand fir (mean 13.8, SE .4 days; *t*(124.3) = 2.45, *p* = .01), but the opposite was true for SNPV (mean 14, SE .4 days on Douglas-fir; mean 12.9, SE .8 days on grand fir; *t*(46.3) = 1.22, *p* = .12), which also had higher variation in its speed of kill (Fig. 3a). Our LOO-CV analysis then showed that the best speed-of-kill model includes effects of morphotype and tree species (Table 3, Fig. 3b). The second-best model includes effects of morphotype only (ΔELPD = −9.4, SE = 6.4), while the third-best accounts for tree species only (ΔELPD = −17.1, SE = 6.6), suggesting that morphotype is a better predictor of speed of kill than tree species.

**Table 3:** LOO-CV analysis of speed-of-kill models. As in previous analyses, for each model we show LOO estimates of the expected log pointwise predictive density (ELPD), the differences in ELPD from the best model (ΔELPD), and the standard error of these differences (SE).

| Model | ELPD | ΔELPD | SE |
| --- | --- | --- | --- |
| Morphotype + tree species | -1070.4 | 0.0 | 0.0 |
| Morphotype only | -1079.8 | -9.4 | 6.4 |
| Tree species only | -1087.5 | -17.1 | 6.6 |
| Intercept | -1091.5 | -21.1 | 7.3 |

**Figure 3:**
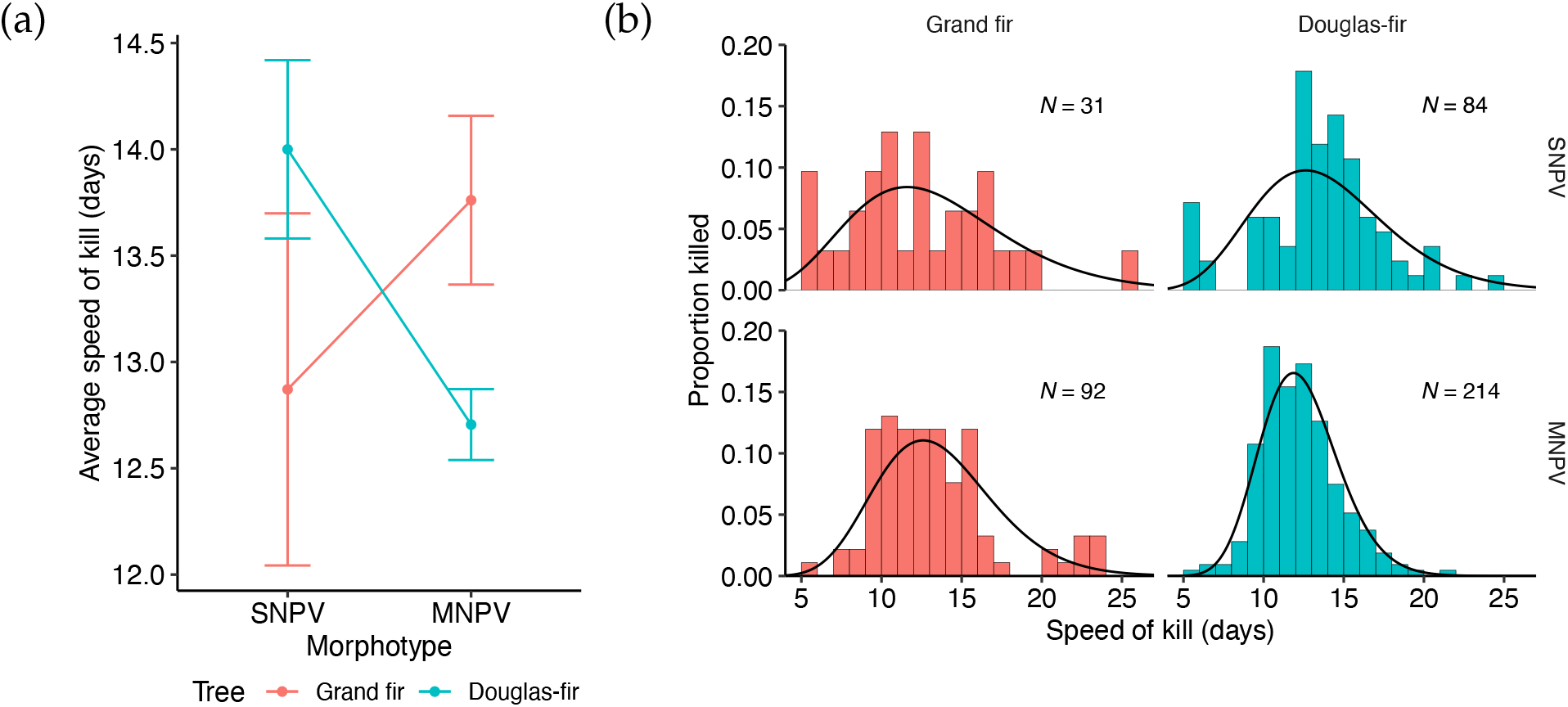
Results of the speed-of-kill experiments. (a) The average number of days from infection to death over all virus-killed larvae, grouped by morphotype and tree species. Points show mean speed of kill and error bars show one standard error of the mean. (b) Speed-of-kill distributions by morphotype and tree species. The best speed-of-kill model, consisting of separate gamma distributions fit to each morphotype and tree species, is shown by the black curves. The sample size *N* is shown for each group.

**Figure 4:**
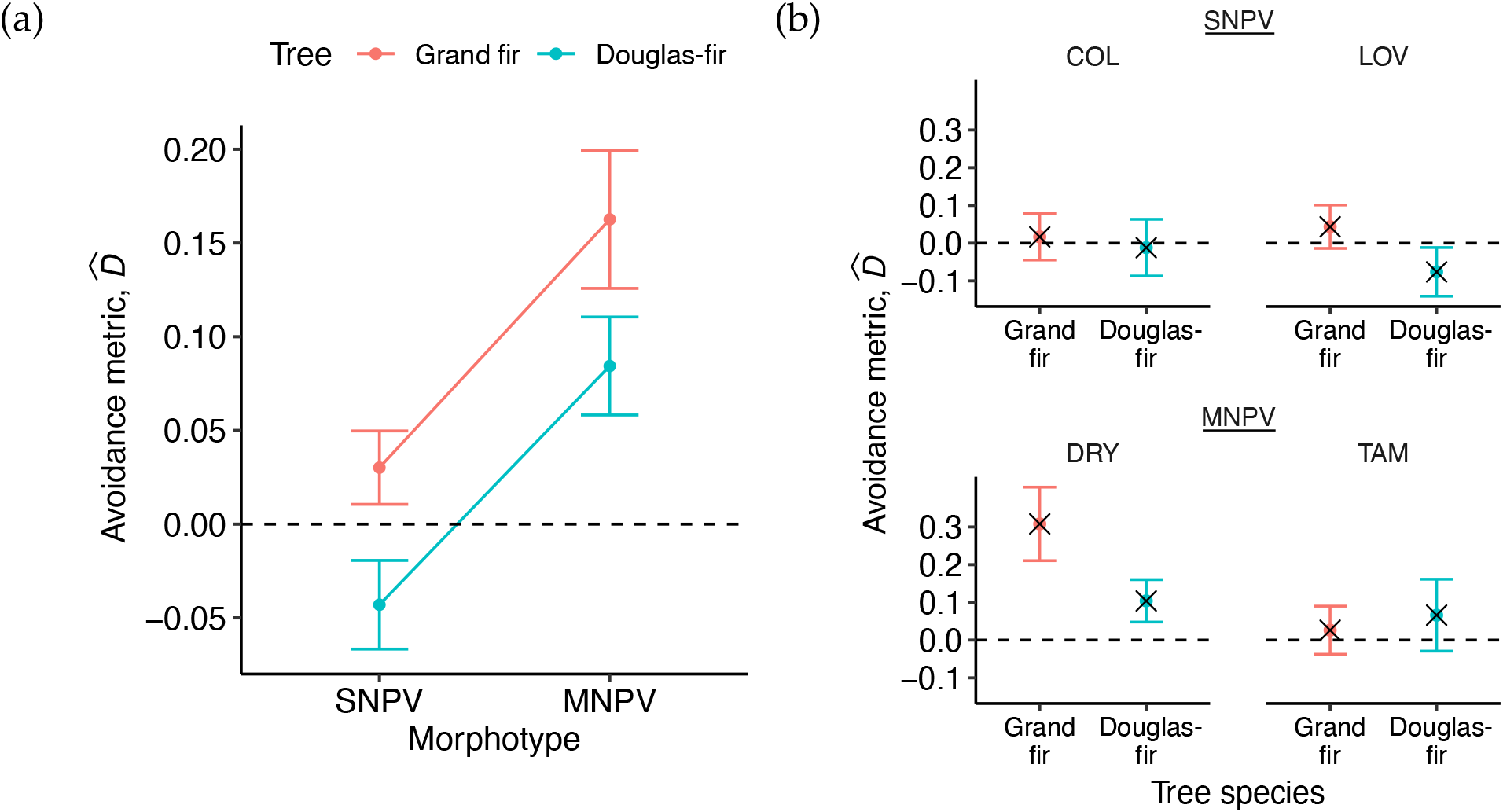
Results of the cadaver avoidance experiments. (a) Average value of the avoidance metric 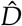 by morphotype and tree species. (b) Average 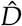 by isolate and tree species, along with the predictions for each group from the best avoidance model, which includes viral presence, morphotype, tree species, and morphotype-tree species interaction (model predictions shown with *×*). In both plots, positive values show cadaver avoidance while negative values show cadaver preference, and the dashed lines at zero represent complete neutrality to cadaver presence. Error bars show one standard error of the mean.

#### Simulation results

Fig. 5a shows the fraction of larvae infected by each viral morphotype in a forest composed of only one tree species or the other. As Fig. 5a shows, if we use our estimated values for each morphotype for both the mean speed of kill and the variance in the speed of kill, then the SNPV morphotype has higher fitness in grand-fir forests, while the MNPV morphotype has higher fitness in Douglas-fir forests. These differences, however, reflect differences in both the mean and the variance. As one would expect for these obligately-lethal pathogens, and as we show in the Online Supplement, faster speed of kill always confers a fitness advantage.

**Figure 5:**
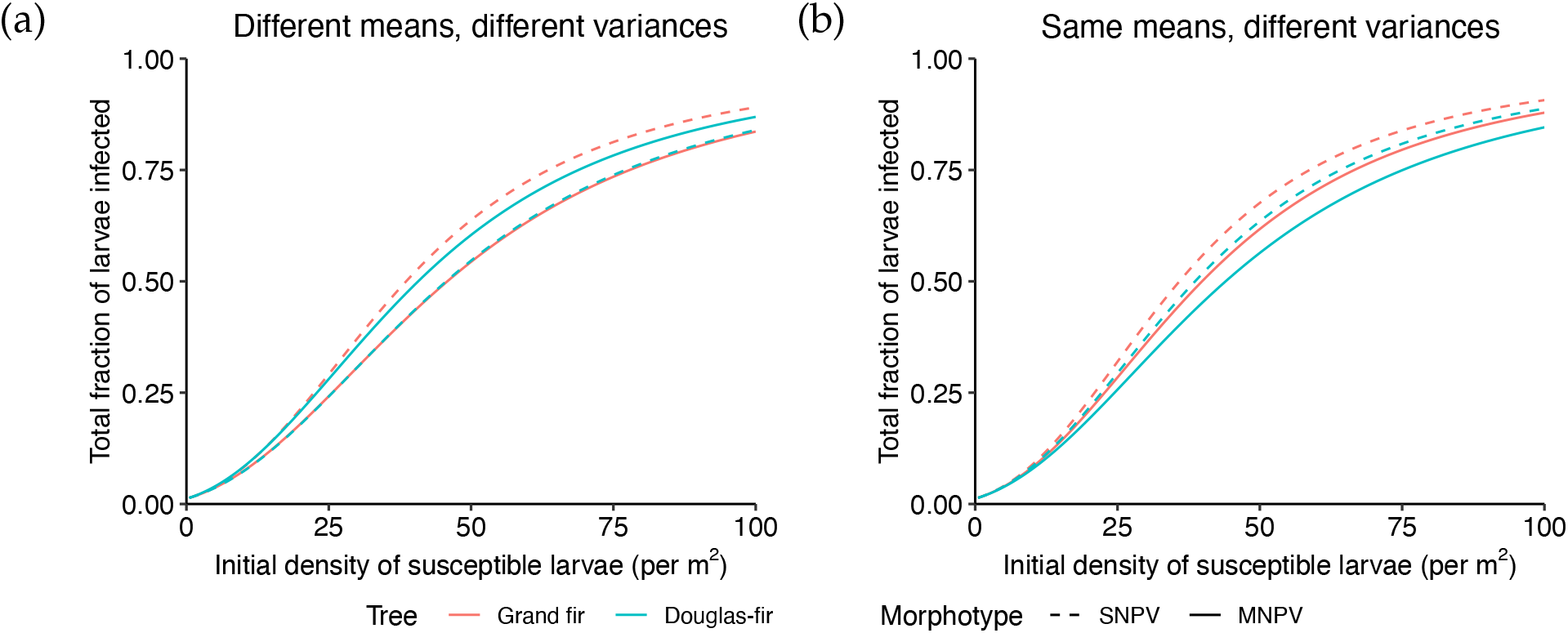
Effects of speed-of-kill distributions on pathogen fitness. For simplicity, here we assume that only one pathogen is present, and that the forest consists of only one tree species. (a) In this case, each morphotype-tree species combination is simulated using the speed-of-kill distribution fit to that specific morphotype and tree species, so that each pathogen-tree species combination has its own mean and variance. (b) In this case, each morphotype-tree species combination is simulated with a speed-of-kill distribution whose variance is specific to that morphotype and tree species but whose mean is equal to the average speed of kill across all morphotypes and tree species. Each morphotype-tree species combination then has its own variance but a different mean. Because the SNPV morphotype has a higher variance in its speed of kill, it has higher fitness on both tree species. In both scenarios, the only parameters differing between the curves are those related to the speed-of-kill distribution.

The effects of differences in the variance in the speed of kill on pathogen fitness, however, are relatively unknown. In Fig. 5b we therefore instead show the case in which each morphotype has the same average speed of kill across morphotypes and tree species, but for which the two morphotypes have the different variances in speed of kill that we estimated from our data. Since the only difference between the two morphotypes in this case is in terms of the variance in the speed of kill, this case isolates the effects of the variance from the effects of the average speed of kill. Fig. 5b then shows that higher variance in speed of kill leads to a fitness advantage, in that the morphotype-tree species combinations with higher variance infect a larger fraction of larvae.

To explain this effect, we note that previous theory has shown that, for obligately-lethal pathogens, a higher fraction of kills early in an epizootic can lead to a higher cumulative fraction infected (Andreasen and Dwyer, 2023). Higher variance in the incubation time therefore appears to confer a fitness advantage because it shifts more infections to earlier in the season, which is essentially the same reason why a shorter speed of kill confers a fitness advantage.

### Summary

Taken as a whole, our data show that the two morphotypes of the baculovirus differ in their transmission phenotypes in multiple ways (Table 4). On grand fir, the higher probability of infection given exposure of the MNPV morphotype confers a fitness advantage, but the shorter average speed, greater variance in the speed of kill and lower cadaver avoidance of the SNPV morphotype give it a compensating advantage. On Douglas-fir, the higher probability of infection given exposure of the MNPV morphotype again confers a fitness advantage, and this advantage is strengthened by the shorter speed of kill of the MNPV morphotype. Meanwhile, on Douglas-fir the SNPV morphotype again has a compensating advantage in terms of the variance of its speed of kill and in terms of its lower cadaver avoidance.

**Table 4:** Summary of fitness advantages by morphotype on each host-tree species, for each component of pathogen fitness. Abbreviations: P(death | exposure) = probability of death given exposure, SOK = speed of kill, and 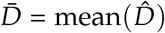 is the average bias-corrected avoidance metric.

| Tree sp. | Component of fitness | SNPV | MNPV | More fit when: | Fitter morphotype |
| --- | --- | --- | --- | --- | --- |
| Grand fir | $\mathbb{P}(\text{death} \mid \text{exposure})$ | 10.5% killed | 26.9% killed | higher | MNPV |
|  | SOK mean | 12.9 days | 13.8 days | lower | SNPV |
|  | SOK variance | 21.2 days<sup>2</sup> | 14.5 days<sup>2</sup> | higher | SNPV |
| | Cadaver avoidance ($\bar{D}$) | 0.0186 | 0.163 | lower | SNPV |
| Douglas-fir | $\mathbb{P}(\text{death} \mid \text{exposure})$ | 23.4% killed | 66.0% killed | higher | MNPV |
|  | SOK mean | 14.0 days | 12.7 days | lower | MNPV |
|  | SOK variance | 14.8 days<sup>2</sup> | 6.0 days<sup>2</sup> | higher | SNPV |
| | Cadaver avoidance ($\bar{D}$) | -0.0284 | 0.0844 | lower | SNPV |

## Discussion

Our results show that four components that are crucial to the fitness of the Douglas-fir tussock moth baculovirus, the probability of infection given exposure, the mean and variance of the speed of kill given infection, and the avoidance of infectious hosts, vary across viral morphotypes and host-tree species. In particular, our results show that the two baculovirus morphotypes have contrasting fitness advantages, and that which morphotype has an advantage with respect to a particular fitness component differs between host trees. Given that the relative frequency of Douglas-fir and grand fir varies strongly across the range of the tussock moth, our results suggest that the coexistence of the two pathogens can be explained by contrasting fitness advantages of the two morphotypes. Notably, of the several fitness components that differ between the MNPV and SNPV morphotypes, only the average speed of kill and the variance in the speed of kill can be easily represented in standard models of pathogen competition. Our work thus suggests that a broader consideration of the ways in which pathogen phenotypes may differ is a key research direction for evolution of virulence theory.

The greater probability of infection given exposure of the MNPV morphotype can likely be attributed to MNPV having about 2.4 times as many virions per occlusion body as SNPV (Hughes, 1979; Martignoni et al., 1971; Rohrmann et al., 1978). The physiological mechanisms by which hostplant foliage interacts with the virus inside the insect, however, are not well understood. Larval diet has been shown to affect multiple steps in baculovirus pathogenesis, including host-plant *p*H effects on the dissolution of the occlusion bodies’ protein coating (Keating et al., 1990), and effects of secondary plant-defensive compounds such as tannins on larval immune responses (Chen et al., 2018; Lee et al., 2006; Trudeau et al., 2001; Washburn et al., 1998, 1996). Tannin concentrations are known to differ in concentration between Douglas and grand fir (Moore et al., 2000), and such variation may help to explain the effects of host-tree foliage in our results. Future work is thus needed to understand the mechanistic basis of pathogen-host diet interactions in the Douglas-fir tussock moth-baculovirus system.

A key feature of our results is that the effect of tree species is significantly larger for MNPV isolates than for SNPV isolates in terms of lethality, implying the presence of a tree species-virus morphotype interaction. Hodgson et al. 2002 similarly observed differential effects of host-tree species on different isolates of the pine beauty moth, *Panolis flammea*. Although Hodgson et al. used isolates obtained from only a single insect and had no information about isolate frequencies in nature, the similarity between our results and theirs suggests that variation in tree species foliage may be of general importance in the maintenance of polymorphism in insect-baculovirus interactions.

A tree-virus interaction is also important in determining the speed of kill of the two morpho-types. These differential effects of tree species on the two morphotypes’ speed and probability of kill suggest that the two morphotypes have evolved distinct life-history strategies on different tree species. The lower cadaver avoidance that larvae showed toward the SNPV morphotype, however, suggests that differences in infection strategies may be complemented by a tradeoff between contact and infectiousness (Lin et al., 2016). The two morphotypes’ infection strategies thus also differ in which transmission steps they specialize on, such that MNPV appears to specialize on successfully infecting and killing its host upon contact, while SNPV specializes on contacting its host in the first place.

Similar genotype-by-environment interactions have been widely invoked for other hostpathogen systems, typically in the context of selection mosaics. Selection mosaics occur when host and pathogen fitnesses vary across environments, so that the outcome of host-pathogen coevolution depends on local environmental conditions (Thompson, 1999). In host-pathogen systems, however, selection-mosaic theory has only ever been applied to plant-pathogen systems (Laine, 2009). Since tussock moth larvae encounter different forest compositions throughout their range, our work implies that viral fitness should vary locally as a result, thus providing a rare example of what appears to be a selection mosaic in an animal host-pathogen system (Dixon, 2024).

Because the tussock moth’s range includes a wide range of forest species compositions and morphotype frequencies, our results may help to provide better predictions of baculovirus dynamics at local geographic scales. Such predictions may make it possible to more effectively use the virus to control the Douglas-fir tussock moth (Martignoni, 1999; Otvos et al., 1987; Scott and Spiegel, 2002). Similarly, our work may make it possible to predict how the dynamics of Douglas-fir tussock moth populations will change over time because of changes in tree composition within the forest (e.g. due to changes in fire regimes) or changes in the relative frequency of morphotypes (e.g. due to the use of a strain of MNPV for tussock moth control; Shepherd et al. 1984). It is also important to acknowledge, however, that our SEIR model is too simple to allow us to project how variation in tree species and morphotype will alter long-term tussock moth dynamics. Models that explicitly allow for spatial variation in tree-species composition of the forest are therefore an important next step (Dixon, 2024).

## Acknowledgments

We thank the Chelsea Miller and Jessica Johnson at the Wenatchee Forestry Sciences Laboratory and William Koval for assistance in data collection both in the lab and in the field. Our work was supported by NSF EEID grant DEB-2109774 and NSF OPUS grant 285 2043796, and by the University of Chicago. A.S.F. was supported by a University of Chicago Ecology and Evolution Fellowship and A.Y.H. was supported by a University of Chicago Biological Sciences Collegiate Division Fellowship.

## Statement of Authorship

G.D. conceived of the project and obtained NSF funding. A.S.F. and A.Y.H. carried out the work with significant help from K.P.D.; among other things, this meant that they figured out how to make the experiments actually work, no small feat. C.P. and G.D. also assisted with the data collection. Data collection was carried out in C.P.’s lab at the Wenatchee Forestry Sciences Laboratory in Wenatchee, Washington. A.S.F. did the bulk of the writing, with editorial contributions from all authors.

## Data and Code Availability

All data and code used to generate figures for this paper can be found at A.Y.H.’s GitHub repository at https://github.com/amyh25/dftm-transmission.

