## Supplementary material for "Effects of Host-Tree Foliage on Polymorphism in an Insect Pathogen": Online Supplement

2. Current address: Department of Ecology and Evolutionary Biology, Princeton University, Princeton, NJ, USA

3. Current address: Computational and Systems Biology, Massachusetts Institute of Technology, Boston, MA, USA

* These authors contributed equally

### Additional Methods

#### *Collection locations and isolate preparation*

Viral isolates were collected from multiple, widely separated locations (Table S1). Of the eight isolates, four are of the SNPV morphotype and the other four MNPV, as identified by transmission electron microscopy. Using a hemocytometer, each isolate was diluted to a viral concentration of 1,800±150 occlusion bodies per $\mu$L, which had been previously established as a relatively low dose for fourth instars. Collection locations and the exact concentrations of each isolate dilution are shown below in Table S1.

| Isolate | Morphotype | Site Name | State | Latitude | Longitude | Concentration |
|---|---|---|---|---|---|---|
| COL* | SNPV | Cheyanne Mountain | CO | 38.73919 | -104.88089 | 1,938 OB/$\mu$L |
| CUB | SNPV | Cub Creek | WA | 47.08265 | -121.23130 | 1,875 OB/$\mu$L |
| LOV* | SNPV | Lovell Valley | ID | 47.30736 | -116.98077 | 1,900 OB/$\mu$L |
| LST | SNPV | Lost River | WA | 48.65235 | -120.50683 | 1,925 OB/$\mu$L |
| DRY* | MNPV | Dry Camp | NM | 35.21516 | -106.40793 | 1,913 OB/$\mu$L |
| KLP | MNPV | Klipchuck Campground | WA | 48.59756 | -120.51301 | 1,663 OB/$\mu$L |
| TAM* | MNPV | Tamarack Flats | ID | 44.387449 | -116.211180 | 1,725 OB/$\mu$L |
| TMB | MNPV | Tussock Moth Biocontrol-1 | CA | 41.96412 | -120.42163 | 1,688 OB/$\mu$L |

Table S1: Collection location and morphotype for each viral isolate used in the infection experiments, as well as the concentrations of the solutions they were diluted down to for our experiments. Starred isolates were also used in the avoidance experiments.

#### *Viral dose consumption in the infection experiments*

As stated in the main text, larvae that did not consume entire needles were discarded. The number of larvae that consumed entire needles dosed with each isolate on foliage of each host tree can be found in Table S2

#### *Foliage collection*

All foliage collected and field experiments conducted were done in areas with very low natural presence of Douglas fir tussock moths or their associated baculovirus. In all experiments, we

| | | Grand fir | | | Douglas fir | | |
|---|---|---|---|---|---|---|---|
| morphotype | isolate | 1 $\mu$L dose | 2 $\mu$L dose | 3 $\mu$L dose | 1 $\mu$L dose | 2 $\mu$L dose | 3 $\mu$L dose |
| SNPV | COL | 22 (36.7%) | 26 (43.3%) | 29 (48.3%) | 32 (53.3%) | 24 (40.0%) | 26 (43.3%) |
| | CUB | 18 (30.0%) | 21 (35.0%) | 20 (33.3%) | 46 (76.7%) | 19 (31.7%) | 18 (30.0%) |
| | LOV | 36 (60.0%) | 8 (13.3%) | 31 (51.7%) | 36 (60.0%) | 46 (76.7%) | 30 (50.0%) |
| | LST | 38 (63.3%) | 19 (31.7%) | 26 (43.3%) | 36 (60.0%) | 26 (43.3%) | 20 (33.3%) |
| MNPV | DRY | 36 (60.0%) | 29 (48.3%) | 37 (61.7%) | 33 (55.0%) | 15 (25.0%) | 29 (48.3%) |
| | KLP | 25 (41.7%) | 47 (78.3%) | 20 (33.3%) | 16 (26.7%) | 26 (43.3%) | 30 (50.0%) |
| | TAM | 25 (41.7%) | 21 (35.0%) | 39 (65.0%) | 45 (75.0%) | 28 (46.7%) | 29 (48.3%) |
| | TMB | 30 (50.0%) | 11 (18.3%) | 22 (36.7%) | 26 (43.3%) | 20 (33.3%) | 27 (45.0%) |

Table S2: Number and proportion of larvae which consumed the entire viral dose in the infection experiments.

collected flush foliage fresh from live trees in the field on the day of the experiment. Foliage would be collected from several trees, to remove confounding effects by individual trees.

#### *Larval rearing*

Douglas fir tussock moth egg masses were collected in May, 2019, from trees in Sage Hen Reservoir, Idaho, then sterilized in 10% formalin before incubating [3]. Larvae were then raised on artificial diet at room temperature with ambient light and humidity [1]. Larvae were reared until the fourth instar, which was chosen to emulate the transmission from neonates (first-instar larvae) infected upon hatching to fourth-instar larvae, a key part of natural tussock moth epizootics [2]. All larval rearing and experiments took place at the Wenatchee Forestry Sciences Laboratory in Wenatchee, Washington.

In the infection experiments, each larva was starved for 24 hours to remove any effects from the artificial diet [4].

#### *Image analysis*

For the image analysis, we used the software ImageJ [8] to manually measure pixel area of fir needles in each Petri dish. In addition, we included a 10 cm section of a ruler in each image, which we measured to convert the pixel area into square centimeters.

### Statistical model descriptions and priors

#### *Dose-response models*

We fit the infection data to four different dose-response models to ascertain whether morphotype and/or tree species were important factors in determining the relationship between viral dose and probability of death given exposure. Each model follows the general logistic-linear form

$$y_n \sim \text{Bin}\left(s_n, \text{logit}^{-1}(\alpha + \beta x_n)\right), \tag{S1}$$

where, $\text{Bin}(k, p)$ is the binomial distribution with $k$ trials and probability $p$ of success; a treatment $n$ that has probability of death given exposure $y_n$, a sample size $s_n$, and a dose $x_n$ in units of occlusion bodies; $\alpha$ and $\beta$ are fitted parameters; and $\text{logit}^{-1} : \mathbb{R} \to (0, 1)$ is the inverse-logit or logistic function defined as $\text{logit}^{-1}(x) = 1/(1 + e^{-x})$. The $\alpha$ and $\beta$ parameters in all models vary by isolate, as well as by tree species for models that have tree species as a factor, and the $\beta$ parameters are always constrained to be positive to avoid unrealistic negative dose responses. The $\alpha$ and $\beta$ parameters are drawn from normal distributions whose means vary by morphotype if that is a factor in the model and by tree species if that is a factor in the model; and whose standard deviations vary only by morphotype if that is a factor in the model but never by tree species. Thus, the isolate-level $\alpha$ and $\beta$ parameters are drawn from a morphotype-level distribution governed by morphotype-level parameters (when morphotype is a factor), to reflect the isolate-morphotype hierarchy that exists naturally. To test the validity of the isolate-morphotype hierarchy in our models, we additionally fit a model with both morphotype and tree species as factors that does not have this hierarchy, by having the parameters governing the distributions for $\alpha$ and $\beta$ existing at the isolate level instead of the morphotype level.

Full mathematical descriptions of these models, as well as the uninformative priors used to fit them, are given in Table S3.

| Morphotype + tree species | Morphotype only |
|---|---|
| $y_n \sim \text{Bin}\left(s_n, \text{logit}^{-1}(\alpha_{i(n),j(n)} + \beta_{i(n),j(n)} x_n)\right)$<br>$\alpha_{i,j} \sim \mathcal{N}(\mu^{\alpha}_{h(i),j}, \sigma^{\alpha}_{h(i)})$<br>$\beta_{i,j} \sim \mathcal{N}(\mu^{\beta}_{h(i),j}, \sigma^{\beta}_{h})$<br>$\mu^{\alpha}_{h,j} \sim \mathcal{N}(0,1)$<br>$\mu^{\beta}_{h,j} \sim \mathcal{N}^{+}(0,1)$<br>$\sigma^{\alpha}_{h} \sim \mathcal{N}^{+}(0,1)$<br>$\sigma^{\beta}_{h} \sim \mathcal{N}^{+}(0,.001)$ | $y_n \sim \text{Bin}\left(s_n, \text{logit}^{-1}(\alpha_{i(n)} + \beta_{i(n)} x_n)\right)$<br>$\alpha_{i} \sim \mathcal{N}(\mu^{\alpha}_{h(i)}, \sigma^{\alpha}_{h(i)})$<br>$\beta_{i} \sim \mathcal{N}(\mu^{\beta}_{h}, \sigma^{\beta}_{h})$<br>$\mu^{\alpha}_{h} \sim \mathcal{N}(0,1)$<br>$\mu^{\beta}_{h} \sim \mathcal{N}^{+}(0,1)$<br>$\sigma^{\alpha}_{h} \sim \mathcal{N}^{+}(0,1)$<br>$\sigma^{\beta}_{h} \sim \mathcal{N}^{+}(0,.001)$ |
| **Tree species only** | **Intercept** |
| $y_n \sim \text{Bin}\left(s_n, \text{logit}^{-1}(\alpha_{i(n),j(n)} + \beta_{i(n),j(n)} x_n)\right)$<br>$\alpha_{i,j} \sim \mathcal{N}(\mu^{\alpha}_{j}, \sigma^{\alpha})$<br>$\beta_{i,j} \sim \mathcal{N}(\mu^{\beta}_{j}, \sigma^{\beta})$<br>$\mu^{\alpha}_{j} \sim \mathcal{N}(0,1)$<br>$\mu^{\beta}_{j} \sim \mathcal{N}^{+}(0,1)$<br>$\sigma^{\alpha} \sim \mathcal{N}^{+}(0,1)$<br>$\sigma^{\beta} \sim \mathcal{N}^{+}(0,.001)$ | $y_n \sim \text{Bin}\left(s_n, \text{logit}^{-1}(\alpha_{i(n)} + \beta_{i(n)} x_n)\right)$<br>$\alpha_{i} \sim \mathcal{N}(\mu^{\alpha}, \sigma^{\alpha})$<br>$\beta_{i} \sim \mathcal{N}(\mu^{\beta}, \sigma^{\beta})$<br>$\mu^{\alpha} \sim \mathcal{N}(0,1)$<br>$\mu^{\beta} \sim \mathcal{N}^{+}(0,1)$<br>$\sigma^{\alpha} \sim \mathcal{N}^{+}(0,1)$<br>$\sigma^{\beta} \sim \mathcal{N}^{+}(0,.001)$ |
| **No hierarchy** | |
| $y_n \sim \text{Bin}\left(s_n, \text{logit}^{-1}(\alpha_{i,j} + \beta_{i,j} x_n)\right)$<br>$\alpha_{i,j} \sim \mathcal{N}(\mu^{\alpha}_{i,j}, \sigma^{\alpha}_{i})$<br>$\beta_{i,j} \sim \mathcal{N}(\mu^{\beta}_{i,j}, \sigma^{\beta}_{i})$<br>$\mu^{\alpha}_{i,j} \sim \mathcal{N}(0,1)$<br>$\mu^{\beta}_{i,j} \sim \mathcal{N}^{+}(0,1)$<br>$\sigma^{\alpha}_{i} \sim \mathcal{N}^{+}(0,1)$<br>$\sigma^{\beta}_{i} \sim \mathcal{N}^{+}(0,1)$ | |

Table S3: The dose-response models. The top four models are those presented in the main text, with the two models that have morphotype as a factor incorporating an isolate-morphotype hierarchy. $\alpha$, $\beta$ are the linear-fit parameters; $\mu_\alpha$, $\mu_\beta$ are their means; and $\sigma_\alpha$, $\sigma_\beta$ are their standard deviations. For a treatment $n$, $y_n$ is the number virus-killed, $s_n$ is the sample size, $x_n$ is the viral dose, $i(n)$ is the isolate, and $j(n)$ is the tree species. All instances of $h$, $i$, or $j$ represent indexing by morphotype, isolate, or tree species, respectively, with $h(i)$ representing the morphotype of isolate $i$. The bottom model is the same as the "Morphotype and tree species" model but without the isolate-morphotype hierarchy (differences colored red). $\mathcal{N}(\mu, \sigma)$ represents the normal distribution with mean $\mu$ and standard deviation $\sigma$, while $\mathcal{N}^{+}(\mu, \sigma)$ is the same distribution but restricted to being positive.

#### *Performance of model with no hierarchy*

Table S4 repeats the main text Table 1, but with the addition of the "No hierarchy" model that is the same as the best model, which has morphotype and tree species as factors, but without the inclusion of the isolate-morphotype hierarchy in the parameter fitting. The "No hierarchy" model does in fact perform worse that the "Morphotype + tree species" model with the hierarchy ($\Delta$ELPD $= -2.6$, SE $= 2.6$). This difference is significant enough that we deem the inclusion of the isolate-morphotype hierarchy in our other models justified.

| Model | ELPD | ΔELPD | SE |
|---|---|---|---|
| Morphotype + tree species | -115.0 | 0.0 | 0.0 |
| Tree species only | -117.6 | -2.6 | 1.7 |
| No hierarchy | -117.7 | -2.6 | 2.6 |
| Morphotype only | -223.6 | -108.6 | 23.0 |
| Intercept | -227.2 | -112.2 | 22.7 |

Table S4: Dose-response models, the LOO estimates of their expected log pointwise predictive densities (ELPD), their differences in ELPD from the best model ($\Delta$ELPD), and the standard errors of these differences (SE). The best model is the one with $\Delta$ELPD $= 0$ and model quality decreases with lower $\Delta$ELPD. The best model is a meaningfully better fit than another model when the magnitude of their $\Delta$ELPD exceeds its SE.

#### *Avoidance models*

Avoidance models quantify the relative importance of viral presence, morphotype, tree species, and morphotype-tree species interaction, the most complex of which incorporates all of these factors:

$$\hat{D}_n = \mu + \gamma_{v(n)} + \rho_{i(n)} + \tau_{j(n)} + \rho_{i(n)}\tau_{j(n)}, \tag{S2}$$

where $\hat{D}_n$ is the avoidance metric (defined in the main text) for Petri dish $n$; $\gamma_{v(n)}$ accounts for viral presence, where $v(n)$ represents whether $n$ is a treatment or control plate, and $\gamma_{\text{control}} = 0$; $\rho_{i(n)}$ accounts for viral morphotype and isolate, such that $i(n)$ is the isolate used in plate $n$ but the standard deviation of $\rho$ varies only by morphotype (accounting for the isolate-morphotype hierarchy), and $\rho_{\text{control}} = 0$; and $\tau_{j(n)}$ accounts for tree species, such that $j(n)$ is the tree species used in plate $n$.

Full mathematical descriptions of these models, as well as the uninformative priors used to fit them, are given in Table S5.

| | |
|---|---|
| Treatment + isolate + tree species + interaction<br>$\hat{D}_n \sim \mathcal{N}(\mu + \gamma_{v(n)} + \rho_{i(n)} + \tau_{j(n)} + \rho_{i(n)}\tau_{j(n)}, \sigma)$<br>$\mu \sim \mathcal{N}(\eta_1, \tau_1) \qquad \gamma_v \sim \mathcal{N}(\eta_2, \tau_2)$<br>$\rho_i \sim \mathcal{N}(\eta_3, \tau_{3,h(i)}) \qquad \tau_i \sim \mathcal{N}(\eta_4, \tau_4)$<br>$\eta_x \sim \mathcal{N}(0, .5)$<br>$\tau_x \sim \text{Gamma}(2, 10)$<br>$\sigma \sim \text{Gamma}(2, 10)$ | Treatment + isolate + tree species<br>$\hat{D}_n \sim \mathcal{N}(\mu + \gamma_{v(n)} + \rho_{i(n)} + \tau_{j(n)}, \sigma)$<br>$\mu \sim \mathcal{N}(\eta_1, \tau_1) \qquad \gamma_v \sim \mathcal{N}(\eta_2, \tau_2)$<br>$\rho_i \sim \mathcal{N}(\eta_3, \tau_{3,h(i)}) \qquad \tau_i \sim \mathcal{N}(\eta_4, \tau_4)$<br>$\eta_x \sim \mathcal{N}(0, .5)$<br>$\tau_x \sim \text{Gamma}(2, 10)$<br>$\sigma \sim \text{Gamma}(2, 10)$ |
| Treatment + isolate<br>$\hat{D}_n \sim \mathcal{N}(\mu + \gamma_{v(n)} + \rho_{i(n)}, \sigma)$<br>$\mu \sim \mathcal{N}(\eta_1, \tau_1) \qquad \gamma_v \sim \mathcal{N}(\eta_2, \tau_2)$<br>$\rho_i \sim \mathcal{N}(\eta_3, \tau_{3,h(i)})$<br>$\eta_x \sim \mathcal{N}(0, .5)$<br>$\tau_x \sim \text{Gamma}(2, 10)$<br>$\sigma \sim \text{Gamma}(2, 10)$ | Treatment + tree species<br>$\hat{D}_n \sim \mathcal{N}(\mu + \gamma_{v(n)} + \tau_{j(n)}, \sigma)$<br>$\mu \sim \mathcal{N}(\eta_1, \tau_1) \qquad \gamma_v \sim \mathcal{N}(\eta_2, \tau_2)$<br>$\tau_i \sim \mathcal{N}(\eta_3, \tau_3)$<br>$\eta_x \sim \mathcal{N}(0, .5)$<br>$\tau_x \sim \text{Gamma}(2, 10)$<br>$\sigma \sim \text{Gamma}(2, 10)$ |
| Tree species only<br>$\hat{D}_n \sim \mathcal{N}(\mu + \tau_{j(n)}, \sigma)$<br>$\mu \sim \mathcal{N}(\eta_1, \tau_1)$<br>$\tau_i \sim \mathcal{N}(\eta_2, \tau_2)$<br>$\eta_x \sim \mathcal{N}(0, .5)$<br>$\tau_x \sim \text{Gamma}(2, 10)$<br>$\sigma \sim \text{Gamma}(2, 10)$ | Treatment only<br>$\hat{D}_n \sim \mathcal{N}(\mu + \gamma_{v(n)}, \sigma)$<br>$\mu \sim \mathcal{N}(\eta_1, \tau_1)$<br>$\gamma_v \sim \mathcal{N}(\eta_2, \tau_2)$<br>$\eta_x \sim \mathcal{N}(0, .5)$<br>$\tau_x \sim \text{Gamma}(2, 10)$<br>$\sigma \sim \text{Gamma}(2, 10)$ |
| Intercept<br>$\hat{D}_n \sim \mathcal{N}(\mu, \sigma)$<br>$\mu \sim \mathcal{N}(\eta, \tau)$<br>$\eta \sim \mathcal{N}(0, .5)$<br>$\tau \sim \text{Gamma}(2, 10)$<br>$\sigma \sim \text{Gamma}(2, 10)$ | |

Table S5: The avoidance models. For a Petri dish $n$, $\hat{D}_n$ is the avoidance metric we measured, $v(n)$ represents whether it is a treatment or control plate, $i(n)$ represents the isolate of the virus used (with $\rho_{\text{control}} = 0$), and $j(n)$ represents the tree species used. All instances of $h$, $i$, or $j$ represent indexing by morphotype, isolate, or tree species, respectively, with $h(i)$ representing the morphotype of isolate $i$. $\mathcal{N}(\mu, \sigma)$ is the normal distribution with mean $\mu$ and standard deviation $\sigma$, and $\text{Gamma}(\alpha, \beta)$ is the gamma distribution with shape $\alpha$ and scale $\beta$.

#### *Speed-of-kill models*

Speed-of-kill models fit gamma distributions to the time from infection onset to death in our infection data, grouping the data and the distributions' parameters by morphotype and tree species, morphotype only, tree species only, or neither. Each model thus follows the general form

$$\tau_n \sim \text{Gamma}(\alpha, \beta), \tag{S3}$$

where $\tau_n$ is the time from infection to death for a successfully infected larva $n$, $\text{Gamma}(\alpha, \beta)$ is the gamma distribution with shape $\alpha$ and scale $\beta$, and $\alpha$ and $\beta$ vary by morphotype and/or tree species. These models do not incorporate the isolate-morphotype hierarchy, since there is not enough data within a single isolate and tree species to specify a gamma distribution well, so we fit parameters only at the morphotype level and never at the isolate level

Full mathematical descriptions of these models, as well as the uninformative priors used to fit them, are given in Table S6.

| | |
|---|---|
| Morphotype + tree species<br>$\tau_n \sim \text{Gamma}(\alpha_{h(n),j(n)}, \beta_{h(n),j(n)})$<br>$\alpha_{h,j} \sim \mathscr{N}^+(0, 10)$<br>$\beta_{h,j} \sim \mathscr{N}^+(0, 10)$ | Morphotype only<br>$\tau_n \sim \text{Gamma}(\alpha_{h(n)}, \beta_{h(n)})$<br>$\alpha_h \sim \mathscr{N}^+(0, 10)$<br>$\beta_h \sim \mathscr{N}^+(0, 10)$ |
| Tree species only<br>$\tau_n \sim \text{Gamma}(\alpha_{j(n)}, \beta_{j(n)})$<br>$\alpha_j \sim \mathscr{N}^+(0, 10)$<br>$\beta_j \sim \mathscr{N}^+(0, 10)$ | Intercept<br>$\tau_n \sim \text{Gamma}(\alpha, \beta)$<br>$\alpha \sim \mathscr{N}^+(0, 10)$<br>$\beta \sim \mathscr{N}^+(0, 10)$ |

Table S6: The speed-of-kill models. For a successfully infected larva $n$, $\tau_n$ is the time from infection to death, $h(n)$ is the viral morphotype used to infect the larva, and $j(n)$ is the tree species on which the larva was fed. All instances of $h$ or $j$ represent indexing by morphotype or tree species, respectively. $\mathscr{N}^+(\mu, \sigma)$ represents the normal distribution with mean $\mu$ and standard deviation $\sigma$, restricted to being positive.

### Epizootic model

To investigate the relative impacts of the mean and variance in speed-of-kill on the dynamics of an epizootic, we simulate an SEIR differential equation model adapated from [6] to explicitly incorporate the gamma speed-of-kill distributions we fit to the infection data. In the main text, we examine a scenario where all morphotype-tree species combinations are simulated with speed-of-kill distributions differing in variance but not mean, and a scenario where the speed-of-kill distributions differ in both variance and mean. Here in the supplement, we also show a scenario where speed-of-kill distributions differ in mean but not variance (Fig. S1c). Simulations differing in speed-of-kill mean/variance use the means/variances taken from our best-fit speed-of-kill distribution model incorporating both morphotype and tree species as factors; simulations using the same speed-of-kill mean/variance use the mean/variance taken from the "Intercept" model that fits just one distribution to all data lumped together.

By examining these three scenarios, we show the effects of changing the speed-of-kill distribution's mean, changing its variance, and changing both the mean and the variance simultaneously. Fig. S1b (Fig. 5b in the main text) shows that higher variance increases the virus's fitness, while Fig. S1a (Fig. 5a in the main text) shows that additionally changing the speed-of-kill distribution's means can complicate these fitness advantages. In Fig. S1c, we isolate the effect of changing only the mean speed-of-kill while keeping the variances the same. From this, we can see a clear advantage for the virus to kill its host faster on average. In reality, however, this may be complicated by the fact that viruses killing their host too fast may not release as many viral occlusion bodies into the environment [5, 7], a factor we do not consider in this model.

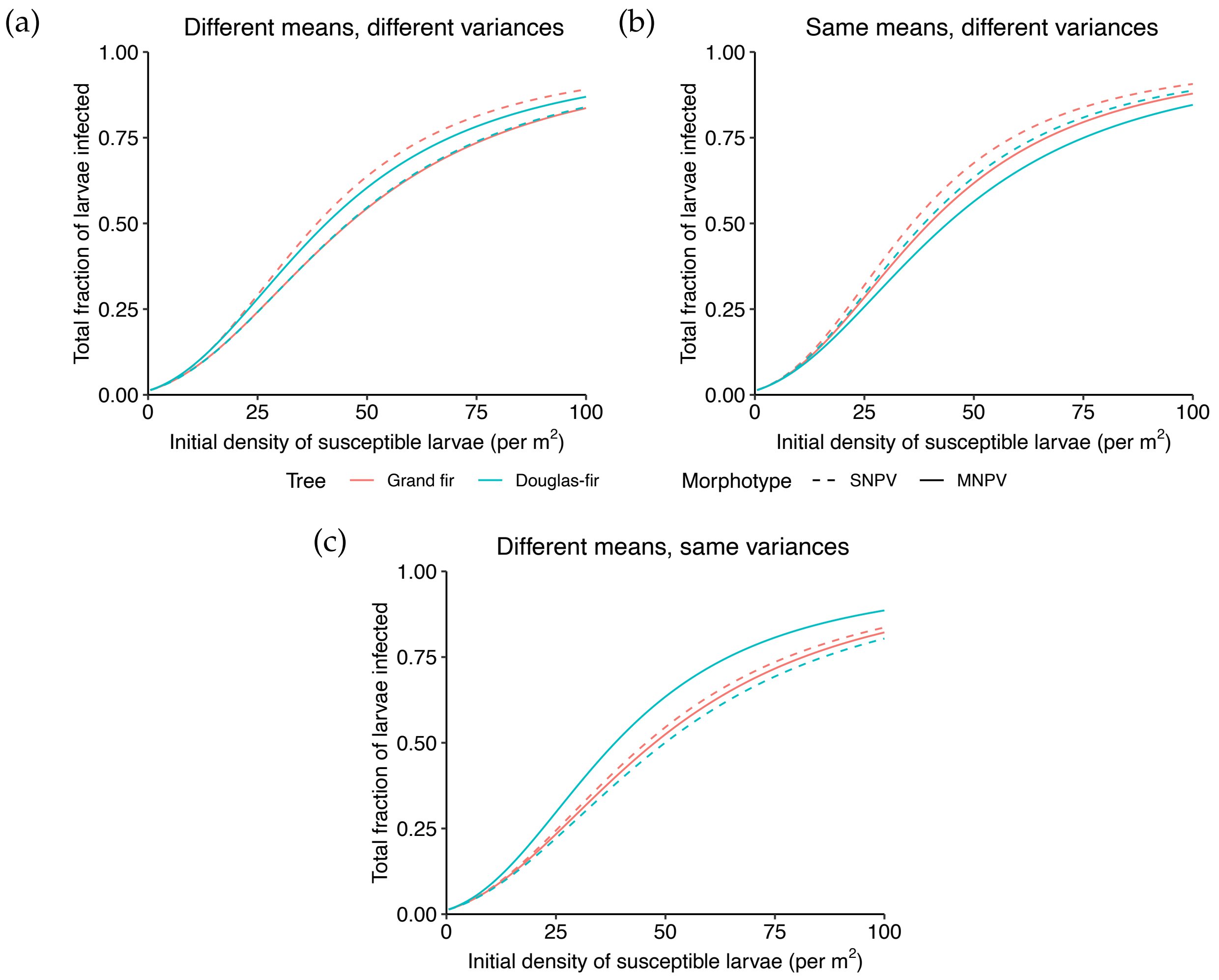


Figure S1: Total fraction of larvae infected in a simulated SEIR epizootic over a range of initial susceptible larva densities, for infections from both morphotypes (line color) and on both tree species (line type). (a) Each morphotype-tree species combination is simulated using the speed-of-kill distribution fit to that specific morphotype and tree species, so that each is simulated with different means and different variances. (b) Each morphotype-tree species combination is simulated with a speed-of-kill distribution whose variance is specific to that morphotype and tree species but whose mean is equal to the average speed of kill across all morphotypes and tree species, so that each is simulated with different variances but the same mean. (c) Each morphotype-tree species combination is simulated with a speed-of-kill distribution whose mean is specific to that morphotype and tree species but whose variance is equal to the variance in speed of kill across all morphotypes and tree species, so that each is simulated with different means but the same variances. In all three plots, the only parameters differing between the curves are those related to the speed-of-kill distribution.

We repeat the model equations here for convenience:

$$\frac{dS}{dt} = -\bar{\nu}SP\left[\frac{S(t)}{S(0)}\right]^{C^2} \tag{S4}$$

$$\frac{dE_1}{dt} = \bar{\nu}SP\left[\frac{S(t)}{S(0)}\right]^{C^2} - m\delta E_1 \tag{S5}$$

$$\frac{dE_i}{dt} = m\delta E_{i-1} - m\delta E_i \qquad (i = 2, \ldots, m) \tag{S6}$$

$$\frac{dP}{dt} = m\delta E_m - \mu P. \tag{S7}$$

Our model differs from that of [6] only in that we disregard stochasticity, and we parameterize $m$ and $\delta$ directly from our speed-of-kill distribution model fits. Specifically, if we wish to model a speed of kill that has mean $\phi$ and variance $\sigma^2$, we set $\delta = 1/\varphi$ and $m = \text{round}(\varphi^2/\sigma^2)$, with $m$ rounded to the nearest integer. Estimates of mean transmission rate $\bar{\nu} = .024$ and coefficient of variation in transmission rate $C = .978$ are taken from [6] from their model fit with all experimental priors, though we set cadaver decay rate $\mu$ equal to 0 to reflect the minimal viral decay that occurs over the course of a single season [9]. The units of $S$, $E_i$, and $P$ are in larvae per m$^2$, and $\bar{\nu}$ has units of per infectious cadaver per m$^2$. The fitness outcome we plot is $1 - S(50)/S(0)$, representing the total cumulative fraction of susceptible larvae that were infected over a 50-day-long epizootic, starting from $P(0) = .01$ and $S(0)$ starting at a value ranging from .5 to 100 susceptible larvae per m$^2$.

in the forest ecosystem. Journal of Invertebrate Pathology 33:57–65.
